# Impact of Noise on Deep Learning-Based Pseudo-Online Gesture Recognition with High-Density EMG

**DOI:** 10.1101/2025.09.08.674890

**Authors:** Mansour Taleshi, Dennis Yeung, Minh Dinh Trong, Francesco Negro, Stéphane Deny, Ivan Vujaklija

## Abstract

Deep neural network (DNN)–based approaches have demonstrated high accuracy in surface electromyography (sEMG)–driven gesture recognition under controlled conditions. However, estimation performance is known to degrade substantially when exposed to perturbations commonly encountered in the real-world. Here, we investigate the impact of typical noises on an autoencoder augmented recurrent neural network gesture estimator driven by engineered features and feature sets. High-density sEMG (HD-sEMG) signals offering rich information particularly suited to DNN-based algorithms were collected from thirteen participants performing wrist movements. Three types of synthetic disturbances were introduced: additive white Gaussian noise (WGN), channel loss, and electrode shift. Results indicate that when using amplitude-based features (specifically, the root mean square value and the mean absolute value), the estimator maintains robust performance under increasing WGN and channel loss, whereas its performance deteriorates markedly with features reflecting signal dynamics and fluctuations, like slope sign changes and zero crossings. Under electrode shift conditions, employing a combined feature set enhances the classifier’s resilience. Importantly, the degree of performance degradation depends on both the type and intensity of the noise. These findings confirm the need for noise-resilient architectures in order to achieve practical, everyday sEMG-driven human-machine interfaces.

**Clinical Relevance:** Quantifying the sensitivity of sEMG-based gesture classifiers to noise can help clinicians tailor electrode placement and training and ultimately decrease user frustration and improve the acceptance of assistive devices.

## I. Introduction

Human-machine interfaces (HMIs) based on surface electromyography (sEMG) signals have emerged as a powerful approach to natural and intuitive control of various devices, ranging from prosthetic limbs [1] to gaming [2] and virtual reality systems [3]. By capturing the electrical activity accompanying muscle contraction, these systems can translate user intent into actionable commands in real-time. Thanks to advances in machine learning methods and computational hardware, many sEMG-based HMIs now achieve high accuracies in controlled settings [4], [5].

The availability of large-scale datasets, particularly those collected using high-density EMG (HD-sEMG) arrays plays an important role in driving HMIs’ advancements. HD-sEMG arrays provide a rich, detailed observation of spatial and temporal information across tens, even hundreds of channels, which in turn allows deep neural networks (DNNs) to learn complex patterns in muscle activity. In fact, large-scale benchmarks have demonstrated that when DNNs are trained on extensive and diverse datasets, EMG-based methods can reliably classify a wide range of gestures with near-real-time response [6], [7]. For instance, large-scale studies such as emg2pose [7] demonstrated that extensive datasets, combined with sophisticated architectures, can dramatically reduce errors in hand pose inference. These developments underscore the notion that neural networks can significantly benefit from the richness and diversity inherent in large-scale data, ultimately leading to more robust and accurate gesture recognition systems.

Despite these promising results in controlled settings, a notable discrepancy remains between laboratory performance and real-world functionality. Namely, high classification accuracy measured offline or under constrained conditions does not always translate into effective functional performance in everyday scenarios [8]. This discrepancy becomes even more evident in more challenging user groups like amputees and other clinical user populations, where system rejection or abandonment rates remain unacceptably high [9]. One key reason is that sEMG signals in daily life are typically subject to noise and perturbations that deviate substantially from the clean recordings of laboratory settings. Factors such as electrode shifts or displacements due to arm movement, sweat-induced impedance changes, environment interference, or corrupted channels can strongly degrade signal quality [10], [11]. As a result, estimation performance in daily use lags behind the near-perfect results reported in controlled experiments.

The study systematically investigates the performance and resilience of DNN models, specifically an autoencoder-augmented recurrent neural network architecture, in the presence of three common perturbations: additive white Gaussian noise (WGN), channel loss, and electrode shift, all of which simulate real-world interactions. Utilizing HD-sEMG sensors across multiple degrees of freedom (DoFs), key global EMG features are extracted to evaluate EMG-based gesture classification resilience against these disturbances. The aim is to identify the sensitivity of different feature sets against signal degradation. By doing so, the study seeks to inform the design of more robust HMIs capable of maintaining high accuracy in realistic, noise-prone conditions.

## II. Methods

### A. Participants

Thirteen able-bodied volunteers (4 females and 9 males of which 12 were right-handed; age range 26–34 years, mean 30) with no known neurological or musculoskeletal disorders took part in this study. Ethical approval was granted by the Aalto University Ethics Committee. All participants provided written informed consent before any experimental procedures began.

### B. Signal Acquisition and Experiment Setup

HD-sEMG and kinematic data were simultaneously collected from each participant’s dominant forearm. Three 8× 8 electrode arrays (ELSCH064NM3, OT Bioelettronica, IT) were placed around the upper half of the forearm to record activity from most muscles contributing to wrist movements. This configuration yielded 192 channels of HD-sEMG data. The reference electrode was positioned near the wrist.

All HD-sEMG signals were sampled at 2048 Hz and digitized via a 16-bit analog-to-digital converter (Quattrocento, OT Bioelettronica, IT). A third-order Butterworth band-pass filter (3–900 Hz) was applied in the hardware. Based on visual inspection, an average of 8*±* 8% of channels were excluded, per subject, due to inconsistent or poor contact during visual inspection (e.g., due to small forearm circumference or skin-electrode interface issues). Prior to electrode placement, participants’ forearms were shaved if necessary, and the skin surface was cleaned with alcohol to further improve the recording quality. All measurements took place in a magnetically shielded room.

Each participant performed three repetitions of six distinct wrist movements including flexion/extension, radial/ulnar deviation, and forearm pronation/supination. These motions were selected to capture a broad range of common daily wrist/forearm actions [12]. Movements were guided by trapezoidal cues displayed on a monitor: participants rested for 2 seconds, followed by 2 seconds each of increasing and decreasing contraction, and finally a steady 10-second contraction at a comfortable level approximating the full range of motion for the targeted motion. Real-time visual feedback of both the desired and actual wrist trajectory derived from upper arm, palm and forearm located wireless inertial measurement units (IMUs) (Xsens Technologies B.V, NL) was provided. This feedback ensure that participants could match the intended profile as closely as possible.

## C. Perturbation Conditions

In order to simulate common signal deteriorations, the recorded “clean” EMGs were exposed to the following three synthetic disturbances

1. *White Gaussian Noise (WGN):* Surface EMG signals are frequently affected by WGN [11], [13], which can arise from factors such as perspiration and degrade the signal-to-noise ratio (SNR). To simulate this, we added varying levels of WGN (10 dB, 9 dB, and up to 1 dB SNR) to the original HD-sEMG data, thereby testing the robustness of neural features under progressively noisier conditions.
2. *Channel Loss:* HD-sEMG data quality is highly sensitive to electrode-skin contact. Poor contact, electrode disconnections, or reduced instrument sensitivity can lead to unusable channels [14]. In many real-world scenarios, losing critical channels might drastically reduce classification performance. To simulate this effect, we set 5%, 10%, and up to 50% of the HD-sEMG channels to zero during the testing phase.
3. *Electrode Shift:* In prosthetics applications, electrode displacement is a recurring issue, particularly when electrodes are inadvertently shifted during everyday activities [15]. Even small shifts can significantly degrade performance if a model has been trained with a fixed electrode configuration. To replicate this, we rotated all electrodes array by one column (1 cm), two columns, and up to 18 columns both laterally and medially (*±*1 cm to *±*18 cm).

### D. EMG Feature Extraction

All signals were segmented into 100 ms windows with a 50% overlap. Within each window, five commonly used global EMG features were extracted (as shown in Figure 1): mean absolute value (MAV), root mean square (RMS), zero crossings (ZC), slope sign change (SSC), and waveform length (WL) [16]. These features capture relevant information in the signals concerning both amplitude and frequency content [16]. The extracted features were then passed to the classifier (section II-E) for assignment to the appropriate class. To evaluate the discriminative power of different feature configurations, the classifier (subsection II-E) was trained with i) each individual feature in isolation, ii) the complete five-feature set, and iii) a compact two-feature subset [RMS, WL]. The RMS + WL pair was selected as a representation because it combines amplitude-based information (RMS) with a measure sensitive to signal dynamics (WL), which results in a balanced yet low-dimensional descriptor.

**Figure 1.**
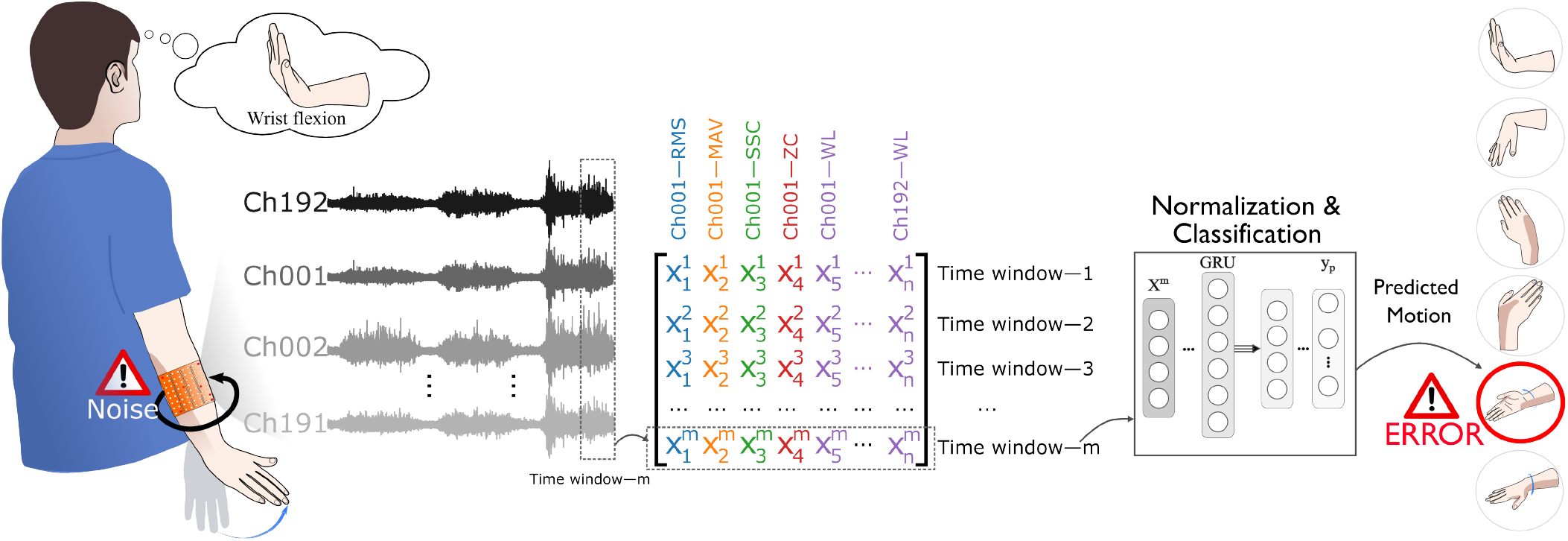
An overview of an EMG-based gesture recognition system. EMG signals from multiple channels (Ch001 to Ch192) are processed to extract features (e.g., RMS, MAV, SSC, ZC, WL), used to classify hand gestures. The system is susceptible to errors when noise perturbs the input signals.

### E. Classification under perturbation conditions

In this study, we employed a DNN or specifically an autoencoder-augmented RNN to classify the motions based on the extracted global features [17], [18]. First, an autoencoder (AE) was trained to learn a compressed latent representation of the input data. The AE comprised a two-layer encoder (each followed by ReLU) and a two-layer decoder (with ReLU and a final Sigmoid), which aimed to reconstruct the original features from the learned embedding. Once trained, the encoder portion was used as input to a gated recurrent unit (GRU) architecture that leveraged the compressed features to capture temporal dependencies [19]. During classification training, a cross-entropy loss was minimized using the AdamW optimizer. During training across multiple epochs, we measured both training and testing accuracies to assess the model’s generalization capability under the 3-fold cross-validation framework.

Following feature extraction and classification, a pseudo-online gesture recognition paradigm was implemented. In this framework, the classifier was first calibrated using EMG features and then evaluated in a simulated real-time setting where perturbations were one-by-one introduced into the EMG signals. Prior to classification, features were normalized using z-score normalization (Figure 1), and a 3-fold cross-validation approach (with two repetitions for training and one for testing) was employed. Classification accuracy (CA) was computed separately for individual features as well as for various combined feature sets.

### F. Statistical Analysis

The Kolgomorov–Smirnov test was first applied to verify data normality. In instances where the data deviated from normality, non-parametric statistical methods were employed. Mean and confidence interval (CI) were calculated, where the CI provides a range in which the true population parameter is estimated to lie. A 95% confidence level was chosen, indicating that there is a 95% probability the true population parameter falls within the calculated CI. All analyses were conducted using Python’s SciPy library.

## III. Results

Figure 2 shows the CA of the DNN classifier across additive WGN, channel loss, and electrode shift perturbations. Each sub-figure shows the performance of the DNN train with one of seven feature sets (the five individual features, a two–feature subset [RMS, WL], and all five combined). Error bars represent the mean *±* 95% CI across participants. Under clean conditions (zero additive WGN or channel loss 0% and no electrode displacement 0 cm), the classifier’s CA was uniformly high (around 80%) for RMS, MAV, WL, their pairwise combination [RMS, WL], and the full combination of all five features ([RMS, MAV, SSC, ZC, WL]). By contrast, SSC and ZC performed at notably lower levels (70%).

**Figure 2.**
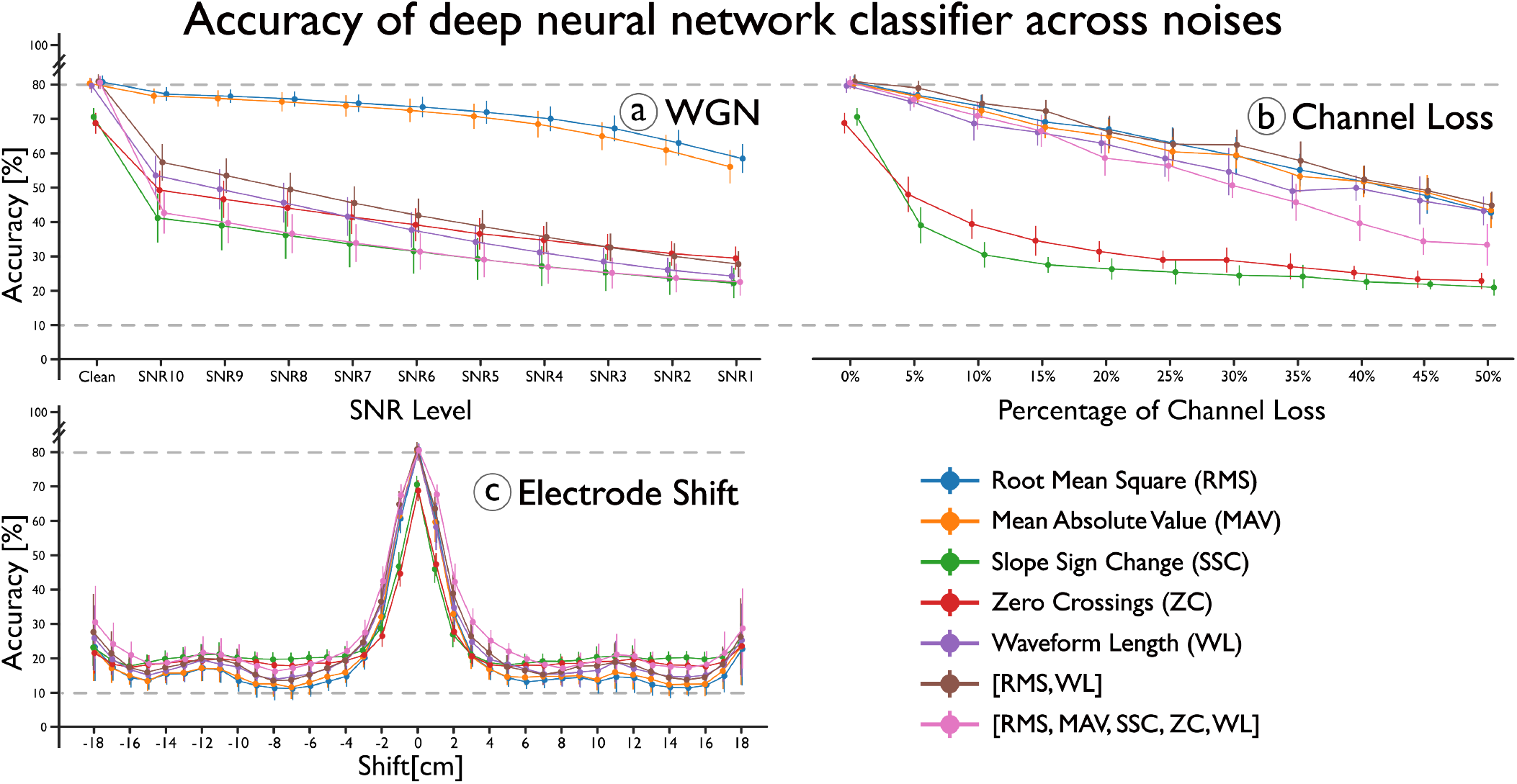
Deep neural network performance under clean (zero additive white Gaussian noise (WGN), 0% channel loss, and 0 cm electrode displacement) and varying noise conditions. Error bars represent the mean *±* 95% CI across participants. (a) classification accuracy (CA) under additive WGN at different SNRs; (b) CA under increasing channel loss (5%–50%); (c) CA under electrode shift (*±*1 cm to *±*18 cm).

As shown by the SNR sweeps from 10 dB down to 1 dB, performance dropped for all feature sets. However, the magnitude and rate of decline varied considerably. RMS and MAV retained the highest CA across moderate to severe noise. At an SNR of 1dB, RMS still maintained 58.57% *±*4.22 and MAV 56.22% *±*4.83, whereas SSC fell to 22.58% *±*4.44 and WL to 24.58%*±* 3.06. Interestingly, combining multiple features under high noise did not always help: the full combination fell to only 22.88% *±*3.93 at 1dB. Even at the relatively mild SNR10 (10dB), RMS and MAV surpassed 76–77% CA, whereas SSC and the full combination dropped to approximately 40–43%. When channels were progressively set to zero (5% through 50%), RMS, MAV, WL, and [RMS, WL] again showed the greatest resilience. At 50% channel loss, RMS maintained 42.81%*±* 4.49, MAV 43.50% *±*5.14, WL 43.30% *±*4.21, and [RMS, WL] 44.93% *±*4.03. SSC and ZC experienced the steepest declines, both dropping to around 21–23% at 50% loss. Notably, the full five-feature combination only reached 33.46%*±* 6.05 at a 50% loss, which was substantially below the single-feature RMS or MAV. Electrode shifts (*±* C1cm up to ±18cm) revealed a different pattern than WGN or channel loss. The full combination displayed the strongest resilience at moderate to large displacements. At *±*1cm shift, the full combination achieved 67.66% *±*3.13, compared to RMS (60.13% *±* 5.03) and MAV (60.63% *±* 5.05). Even at *±*18cm, where all features dropped substantially, the full combination still reached 29.69% *±* 11.12, which was higher than RMS (23.11% *±* 10.24) and MAV (23.30% *±* 10.35).

## IV. Discussion

This study examined the sensitivity of a DNN–based gesture recognition to a common representative of real-world noise simulated through additive WGN, channel loss, and electrode shift signal perturbations. The result showed that RMS and MAV retain higher classification accuracy under moderate to severe WGN and increasing channel loss, while SSC and ZC degrade more rapidly, likely due to their greater sensitivity to rapid amplitude and frequency fluctuations in the signal. Following electrode shift, the combined feature set performs better than any single feature alone, suggesting that spatial realignment challenges are alleviated by more diverse feature representations. Among the three perturbation types, electrode shift leads to the most pronounced performance drop. This impact highlights the importance of stable sensor placement for reliable real-world use. One possible explanation is that simple amplitude-based features (RMS, MAV) remain robust when electrodes stay in the same location, but once the spatial configuration changes, the synergy of multiple features is needed to preserve discriminatory power.

A major limitation of this study is that it focuses on simulated well-defined noise sources in able-bodied individuals and the lack of user-in-the-loop adaptation; in practice, unknown or more complex noise can arise, and user populations may differ substantially. Additionally, this work relies on explainable, hand-crafted features (e.g., RMS, MAV, WL) rather than directly on the raw HD-sEMG signals. While feature engineering affords interpretability and computational efficiency, it necessarily constrains the model to pre-selected signal characteristics. End-to-end approaches such as sliding-window convolutional neural networks or other deep architectures trained on raw EMG may discover complementary representations and potentially yield better robustness under noise.

While it is well known that noise degrades EMG-based gesture recognition performance, our study provides a detailed scaling of this effect. We demonstrated that even a small channel loss can be highly impactful–especially when using certain combinations of features–while amplitude-based features (e.g., RMS and MAV) consistently yield higher accuracy under additive white Gaussian noise, with performance closely tied to the SNR level. These results emphasize that not all features are affected equally by noise and that careful selection and combination of features can mitigate performance loss to some extent. Accordingly, depending on the expected usage, it may be necessary to adjust network design and feature selection in accordance with the environment and intended application.

Ultimately, although the exemplary DNN implementation presented here performs reasonably under clean conditions, these results underscore that the model optimization alone cannot effectively address all perturbations. It is important to note that this implementation represents only one approach and alternative architectures might exhibit different sensitivities to noise and perturbations. Moreover, while this study did not incorporate actual user-in-the-loop scenarios, integrating adaptive user feedback may alter the outcomes. Developing adaptive methods that continuously learn and recalibrate might offer a way to bridge the gap between high accuracies in clean settings and reliable daily use. This can be achieved through co-adaptive strategies [20], user-in-the-loop training [11], or self-supervised approaches, such as the Barlow Twins paradigm [21], which naturally reduces redundancy in high-dimensional embeddings and may better tolerate electrode displacement or unseen noise. Finally, incorporating simulated or real-world perturbations into the training process may further improve the model’s robustness in practical settings [22], [23].

## Acknowledgment

The authors are grateful to the Aalto Science-IT project for providing computational resources. This work was supported in part by the Academy of Finland under Grant 333149 (Hi-Fi BiNDIng) for M.T. and I.V.-D.Y., and under Grant 3357590 for S.D. Additionally, support was provided by the Jenny and Antti Wihuri Foundation under Grant 00230401 for M.T., the European Research Council (ERC) through the Consolidator Grant INcEPTION (Contract No. 101045605) for F.N., and the Helsinki Institute for Information Technology (HIIT) under Contract No. 9125064HIIT for M.D.T.

## Notes

https://m-taleshi.github.io/publications/

### Competing Interest Statement

The authors have declared no competing interest.

